# A systematic, label-free method for identifying RNA-associated proteins *in vivo* provides insights into vertebrate ciliary beating

**DOI:** 10.1101/2020.02.26.966754

**Authors:** Kevin Drew, Chanjae Lee, Rachael M. Cox, Vy Dang, Caitlin C. Devitt, Ophelia Papoulas, Ryan L. Huizar, Edward M. Marcotte, John B. Wallingford

## Abstract

Cell-type specific RNA-associated proteins (RAPs) are essential for development and homeostasis in animals. Despite a massive recent effort to systematically identify RAPs, we currently have few comprehensive rosters of cell-type specific RAPs in vertebrate tissues. Here, we demonstrate the feasibility of determining the RNA-interacting proteome of a defined vertebrate embryonic tissue using DIF-FRAC, a systematic and universal (*i.e*., label-free) method. Application of DIF-FRAC to cultured tissue explants of *Xenopus* mucociliary epithelium identified dozens of known RAPs as expected, but also several novel RAPs, including proteins related to assembly of the mitotic spindle and regulation of ciliary beating. In particular, we show that the inner dynein arm tether Cfap44 is an RNA-associated protein that localizes not only to axonemes, but also to liquid-like organelles in the cytoplasm called DynAPs. This result led us to discover that DynAPs are generally enriched for RNA. Together, these data provide a useful resource for a deeper understanding of mucociliary epithelia and demonstrate that DIF-FRAC will be broadly applicable for systematic identification of RAPs from embryonic tissues.

## Introduction

RNA-associated proteins (RAPs) play diverse roles in normal cellular physiology and their disruption is linked to diverse pathologies (Castello et al., 2013; Gerstberger et al., 2014; Hentze et al., 2018; Quinn and Chang, 2016; Wickramasinghe and Venkitaraman, 2016). Encompassing both direct RNA-binding proteins and indirect interactors, the RAP universe is expanding rapidly. Indeed, there is now a global effort to systematically identify and characterize RAPs, and this effort continues to provide new biological insights as well as invaluable resources for hypothesis generation (e.g. (Baltz et al., 2012; Bao et al., 2018; Brannan et al., 2016; Castello et al., 2016; Caudron-Herger et al., 2019; He et al., 2016; Huang et al., 2018; Mallam et al., 2019; Queiroz et al., 2019; Treiber et al., 2017; Trendel et al., 2019)).

However, while this effort has been focused largely on a narrow selection of cultured cell types, a wide array of *cell-type specific* RAPs are known to be essential for normal development and homeostasis. For example, RAPs control the specific transport and localization of mRNAs in cells ranging from totipotent fertilized eggs to highly differentiated cells such as neurons (Holt and Bullock, 2009; Medioni et al., 2012; Sahoo et al., 2018). Moreover, mammalian ribosomal proteins are now thought to play cell-type specific roles, for example in the translation of developmental regulators such as Hox genes during mouse development (Kondrashov et al., 2011; Xue and Barna, 2012), in development of the spleen (Bolze et al., 2013), and in hematopoiesis (Choesmel et al., 2007). Recent work in *C. elegans* has argued against specialized ribosomes (Cenik et al., 2019; Haag and Dinman, 2019), but *C. elegans* nonetheless displays a curious non-cell-autonomous requirement for ribosomes in growth control (Artiles et al., 2019; Cenik et al., 2019).

At the same time, RNA-associated proteins have also emerged as critical factors for the assembly of liquid-like organelles, including not only ubiquitous organelles such as nuceloli and stress granules, but also cell-type specific organelles such as Cajal bodies (Banani et al., 2017; Lin et al., 2015; Mittag and Parker, 2018; Sawyer et al., 2016). In a recent study, we described novel liquid-like organelles called DynAPs that are specific to multiciliated cells, where they control the assembly of dynein motors that drive ciliary beating (Huizar et al., 2018). DynAPs are present in motile ciliated cells of *Xenopus*, zebrafish, and mammals, and their disruption is associated with human motile ciliopathy (Horani et al., 2018; Huizar et al., 2018; Li et al., 2017; Mali et al., 2018). Thus, understanding the role of RNA-associated proteins in DynAPs is an important specific challenge, while development of methods for systematic identification of RAPs from animal tissues will be broadly useful.

Here, we demonstrate the feasibility of determining the RNA-interacting proteome of a defined vertebrate embryonic tissue using DIF-FRAC, a systematic and universal (*i.e.*, label-free) method (Mallam et al., 2019). Application of DIF-FRAC to cultured tissue explants of *Xenopus* mucociliary epithelium identified over 380 RNA-associated proteins, including dozens of known RAPs, but also several novel RAPs. Many of the novel RAPs play roles in assembly of the mitotic spindle and the regulation of ciliary beating. These data provide a useful resource for a deeper understanding of mucociliary epithelia and demonstrate that DIF-FRAC will be broadly applicable for systematic identification of RNA-associated proteins in both model and non-model organisms.

## Results

We recently developed a novel method for systematically identifying RNA-protein interactions based on differential fractionation (DIF-FRAC)(Fig. 1)(Mallam et al., 2019). In DIF-FRAC, a protein lysate from theoretically any native or non-denatured sample is split into two; one is treated with RNAse and the other serves as a control; each sample is then independently subjected to size-exclusion chromatography (SEC) and the contents of each fraction are independently quantified by mass spectrometry. Shifts in elution profiles reveal alterations to protein complexes in the presence or absence of RNA, and a novel statistical framework is used to quantify these changes and thereby identify RNA-associated proteins (Fig. 1A).

**Figure 1:**
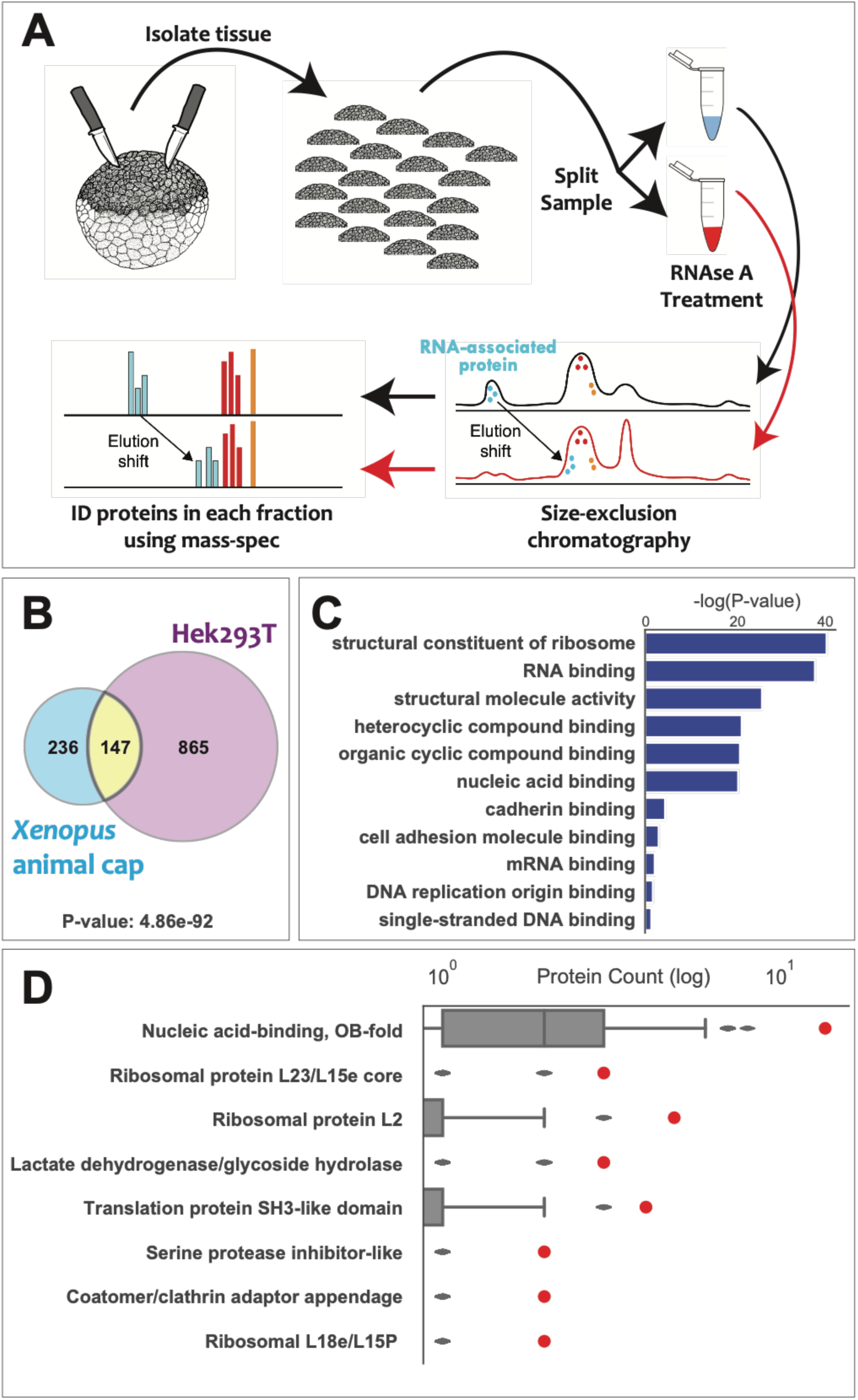
Differential Fractionation (DIF-FRAC) identifies RNA-associated proteins in a mucociliary epithelium. (A) Experimental workflow of the RNAse DIF-FRAC experiment on *Xenopus* animal cap explants. (B) Venn diagram displaying overlap of RNA-associated proteins in *Xenopus* animal caps with previously published data from HEK293T cells (Mallam et al., 2019). The *p*-value represents the probability of overlap based on chance using the hypergeometric test. (C) Gene ontology molecular function enrichment analysis of significant DIF-FRAC hits from *Xenopus* (D) Interpro superfamily enrichment analysis. Red dots represent the count of identified RNA-associated proteins annotated with individual Interpro superfamilies. Gray box plots represent the background distribution of proteins annotated with individual Interpro superfamilies.

To test the efficacy of DIF-FRAC on embryonic tissue, we used the pluripotent ectoderm from gastrula stage *Xenopus* embryos (so-called “animal caps”), which can be readily transformed into a wide array of organs and tissues, including a motile ciliated epithelium (Ariizumi et al., 2009; Walentek and Quigley, 2017; Werner and Mitchell, 2012). This tissue can be obtained in abundance, as we and others have demonstrated in large-scale genomic studies of ciliogenesis and cilia function (Chung et al., 2014; Ma et al., 2014; Quigley and Kintner, 2017). As outlined in Figure 1A, we collected ∼3,000 micro-dissected animal caps and cultured them to stage 23, when motile cilia have been assembled and begin to beat. We performed DIF-FRAC using ∼80 SEC fractions each for control and RNAse treated samples.

Due to the allotetraploid nature of the *Xenopus laevis* genome (Session et al., 2016), multiple copies of most genes exist in the genome, hindering accurate identification of proteins by mass spectrometry. Previous approaches used a transcriptome-based reference proteome (Wühr et al., 2014), but this approach is limited to proteins whose transcripts were identified in the employed mRNA sequencing datasets and therefore under-represent proteins with highly restricted expression, for example those related to ciliary motility.

To overcome these limitations, we applied an orthology-based approach that we recently developed for proteomic comparisons across polyploid plant species (McWhite et al., 2019), in which we collapse highly related protein sequences into EggNOG vertebrate orthology groups (Huerta-Cepas et al., 2016)(Supp. Figure 1, see methods). Using this approach, we substantially increased the number of uniquely assigned mass spectra, as well as the number of identified orthology groups compared to the standard *Xenopus laevis* reference proteome (Supp. Table 1). Peptide spectral matches (PSMs) for each orthology group were then used to construct elution profiles (*i.e.*, abundance of protein across fractions) and elution profiles were compared across control and RNAse treatments.

To identify RNA-associated proteins in DIF-FRAC data, we use a computational framework to identify statistically significant changes in a protein’s elution behavior (the “DIF-FRAC score”) which allows us to assign *p*-values by comparing each protein’s DIF-FRAC score to an abundance-controlled background distribution of DIF-FRAC scores from non-RNA-associated proteins. (A fuller explanation can be found in the Methods section and in our published work (Mallam et al., 2019).) This *p*-value takes into account the observed abundance as well as the size of the elution profile shift. This statistical framework provides a ranked confidence score for each protein, and previous work suggested that 0.05 is a reasonable initial threshold (Mallam et al., 2019).

Using this approach, we identified over 380 RAPs in *Xenopus* animal caps, and this set displayed significant overlap with the set previously identified in Hek293T cells (Fig. 1B)(Supp. Table 2). Our significant DIF-FRAC hits from *Xenopus* were also highly enriched for GO annotations associated with RNA and RNA binding (Fig. 1C). Finally, an analysis of protein superfamilies using the Interpro database (Hunter et al., 2009) revealed that proteins with significant DIF-FRAC scores were also highly enriched in protein domains associated with RNA-binding, nucleic acid binding, or the ribosome (Fig. 1C).

To complement these systematic analyses, we manually curated our high-scoring hits from *Xenopus* DIF-FRAC. We found, for example, that of the 60 highest scoring ribosomal proteins, 52 displayed DIF-FRAC scores <0.05, with 46 scoring p<0.01 (Fig. 2A-D). We hasten to note, however, that any specific threshold should be interpreted with care. Indeed, additional ribosomal proteins scored above this threshold in our dataset despite having a generally consistent elution profile shift similar to other ribosomal proteins (Fig. 2E).

**Figure 2:**
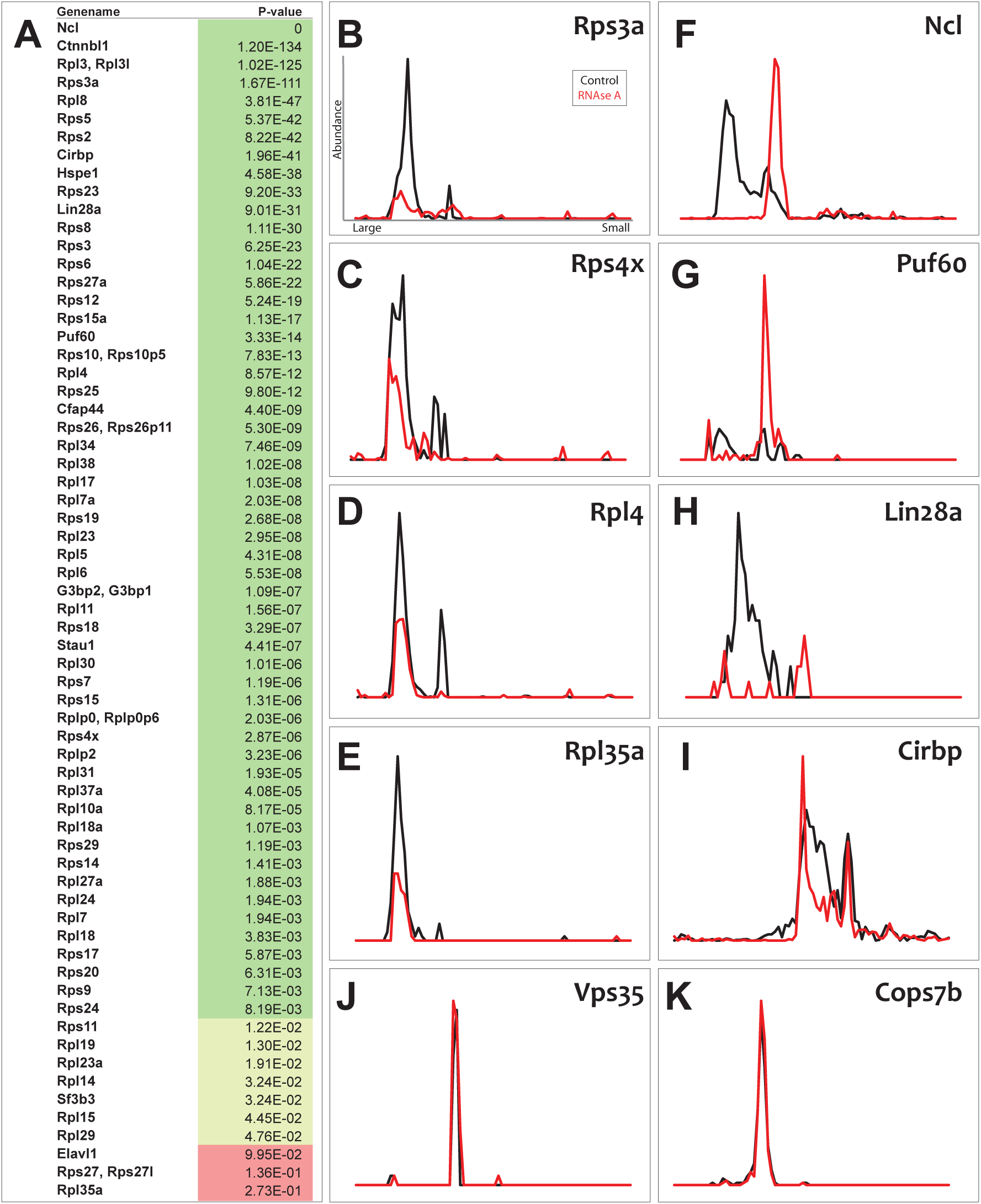
Individual DIF-FRAC elution profiles show distinct changes consistent with RNAse sensitivity. (A) Table of DIF-FRAC calculated *p*-values for ribosomal proteins and selected others. (B-D) Individual profiles for ribosomal subunits. X-axis represents SEC fractions from larger molecular weight to smaller. Y-axis represents observed abundance in MS by unique peptide spectral matches normalized to the highest value for that protein. (E) Ribosomal subunit, Rpl35a, had a *p*-value above the 0.05 cutoff (A), but its elution profile shows consistent behavior with other ribosomal subunits. (F) Known RNA-binding protein, Nucleolin, shows shift in molecular weight. (G) Known RNA-binding protein, Puf60, shows increased observed abundance. (H-I) Profiles of RNA-binding proteins with known roles in *Xenopus* development. (J-K) Elution profiles of negative controls do not change.

Other significant hits also included known components of universal RNA processing machines, such as nucleoli (Ncl), the spliceosome (Sf3b3, Puf60), and stress granules (G3bp2)(Fig. 2F-G, Supp. Table 2). More importantly, we identified several RNA-binding proteins with known roles in early *Xenopus* embryos, such as Lin28a, Staufen1, and Cirbp (Faas et al., 2013; Peng et al., 2006; van Venrooy et al., 2008; Yoon and Mowry, 2004)(Fig. 2H-I). By contrast, known negative controls such as Vps35 and Cop9 signalosome subunits showed no elution shift after RNAse treatments (Fig. 2J, K). An independent biological replicate of our DIF-FRAC experiment yielded similar results, confirming the robustness of the method (Supp. Fig. 2, Supp. Table 3).

DIF-FRAC in cultured cells resulted in distinct classes of altered elution profiles for distinct RAPs (Mallam et al., 2019), and we observed similar trends in our data from *Xenopus*. For example, Ncl displayed a clear mobility shift after RNAse treatment, but the protein continued to elute in defined peaks (Fig. 2F), suggesting that it is stable in the absence of RNA. On the other hand, for ribosomal proteins and Lin28a, overall observed abundance was drastically reduced; no clear peaks were observed after RNAse treatment (Fig. 2B-E, H), indicating reduced protein stability in the absence of RNA. Finally, the splicing factor, Puf60, displayed a clear *increase* in observed abundance (Fig. 2G), likely because it is insoluble when complexed with RNA. Thus, both systematic analysis and manual curation indicate that DIF-FRAC effectively identified RAPs in *Xenopus*.

Importantly, our list of high scoring proteins also contained several proteins not previously known to associate with RNA, providing new insights. For example, our significant DIF-FRAC hits were enriched with the “Lactate dehydrogenase/glycoside hydrolase” superfamily (Fig. 1D), consistent with recent data suggesting frequent interaction of metabolic enzymes with RNA and suggesting ‘moonlighting’ functions (Castello et al., 2015). The “Coatomer/clathrin adaptor appendage” superfamily was also enriched in our significant DIF-FRAC hits (Fig. 1D), including for example AP-2 complex sub-units (Supp. Fig. 3B-C). This result is consistent with reports of RNA interactions with the related COPI complex (Todd et al., 2013).

**Figure 3:**
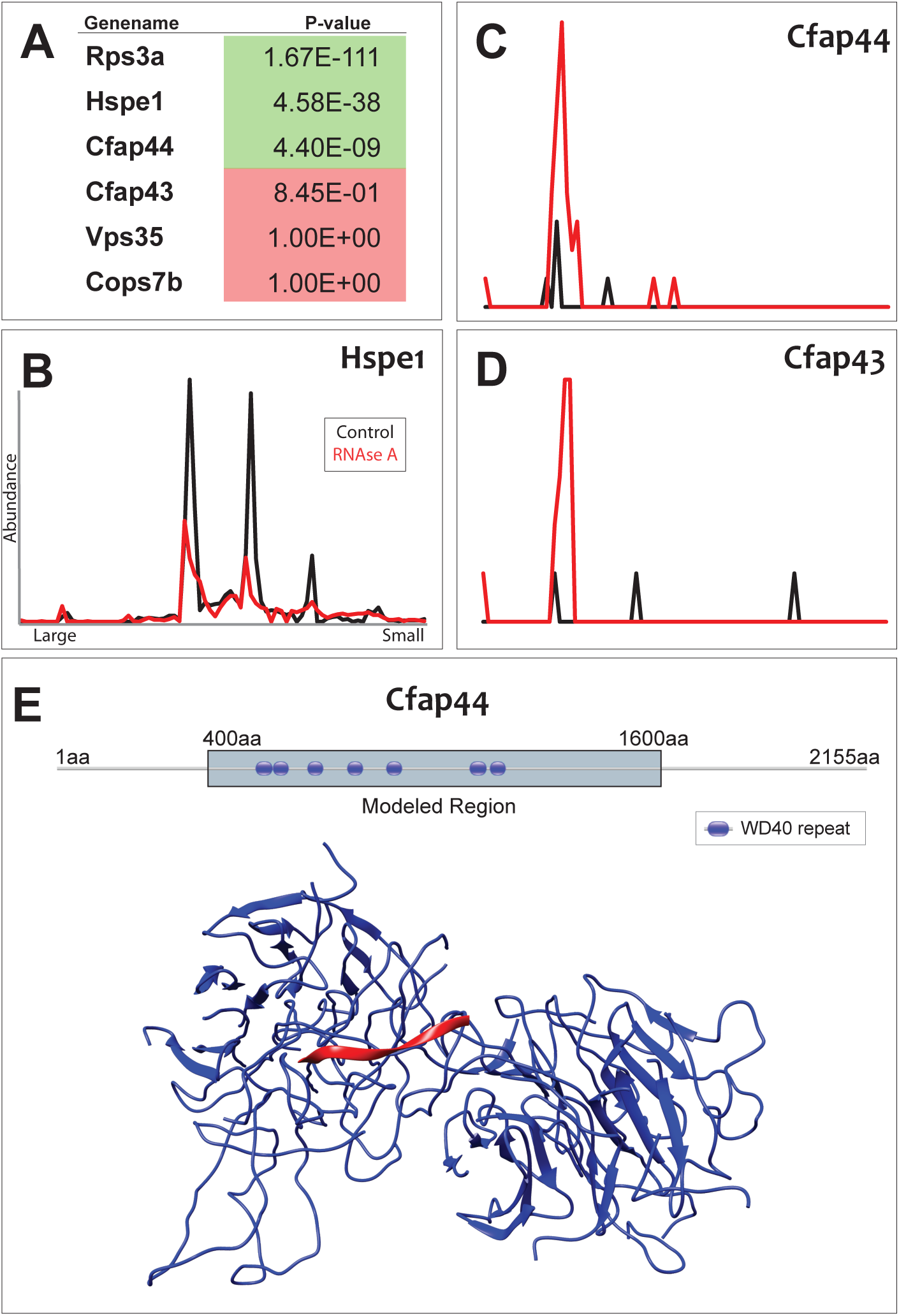
DIF-FRAC identifies a ciliopathy protein as RNA associated. (A) Table of DIF-FRAC calculated *p*-values for selected motile cilia-related proteins; Rps3A, Vps35 and Cops7b serve as positive and negative controls. (B) Elution profile of Hspe1 shows loss of observed abundance. (C) Elution profile of the inner arm dynein tethering protein Cfap44 shows a gain of observed abundance. (D) Elution profile of Cfap43 shows similar behavior to Cfap44. (E) Homology model of Cfap44 WD-40 domains (blue) with an RNA molecule (red) is modeled from Gemin5 crystal structure (PDB ID: 5GXH).

Our dataset also contained a number of microtubule-associated proteins not previously known to be RNA-associated. Among these, Ccdc124/Lso2 was of interest because it is linked to both ribosomes and the mitotic spindle (Telkoparan et al., 2013; Wang et al., 2018). Several recent studies have explored spindle-associated RNAs (Blower et al., 2007; Jambhekar et al., 2014; Sharp et al., 2011), so it is notable that several additional spindle-associated proteins were identified here as RAPS. In both replicates of *Xenopus* DIF-FRAC, we identified Kif11/Eg2, Kif15, Map1s, Eml1, Eml2, and Map7d3 (Supp. Tables 2, 3). These findings are of special interest in the context of multiciliated cells, since much of spindle machinery is shared with motile and primary cilia (Bernabé-Rubio et al., 2016; Smith et al., 2011).

We also identified several novel RAPs that also have potential or known roles in ciliary beating. These include the axonemal dynein subunit Dnah11, the radial spoke protein Rsph1, and Hspe1, a heat shock family chaperone known to interact with the axonemal protein Spag16, which is also present in liquid-like nuclear speckles (Nagarkatti-Gude et al., 2011; Zhang et al., 2008) (Fig. 3A-B). The most interesting novel RAP identified here was Cfap44, encoded by a human ciliopathy gene and required for cilia motility in diverse organisms (Coutton et al., 2018; Fu et al., 2018; Kubo et al., 2018; Tang et al., 2017). The RNA-dependent change in elution profile for Cfap44 was unexpected, but we observed this elution shift for Cfap44 in both biological replicates (Supp. Fig. 2F). In both samples, Cfap44 displayed the distinct “gain in observed abundance” pattern described above for Puf60, which we interpret as a gain in protein solubility upon RNA degradation (Mallam et al., 2019).

In our previous work, incorporation of prior knowledge of protein-protein interactions allowed for identification of RAPs that may not qualify as statistically significant by their elution profiles alone (Mallam et al., 2019). This was also the case in our *Xenopus* DIF-FRAC (e.g. Rpl35a, Fig. 2A, E). This principle led us to then examine the elution profile of Cfap43, a known interaction partner of Cfap44 (Urbanska et al., 2018). Importantly, despite a relatively high DIF-FRAC *p*-value (Fig. 3A), Cfap43 displayed a clear shift upon RNAse treatment that was strikingly similar to that of Cfap44 (Fig. 3D), suggesting that these two proteins may interact with RNA as a complex.

As an independent assessment as a candidate RAP, we used homology modelling to explore the structure of Cfap44. Using both pGenTHREADER and HHPred, we found that Cfap44 has similarity to the known RNA-associated protein Gemin5 (PDB ID: 5GXH)(Battle et al., 2006; Xu et al., 2016; Yong et al., 2010). We further found a WD40 domain between amino acids 400 and 1600 in Cfap44 (Fig. 3E) and using the existing co-crystal structure of Gemin5 bound to RNA (Xu et al., 2016), we were able to model an RNA molecule with our Cfap44 structural model (Fig. 3E).

These results prompted us to consider what role an interaction with RNA may play in Cfap44 function. We observed strong labelling of MCC axonemes with Cfap44-GFP (Fig. 4A, a’), consistent with reports from other organisms (Coutton et al., 2018; Fu et al., 2018; Kubo et al., 2018; Urbanska et al., 2018). However, because RNA is not thought to act in axonemes, we re-examined the localization of Cfap44 in more detail. Strikingly, Cfap44-GFP also localized strongly to foci in the cytoplasm of *Xenopus* MCCs (Fig. 4B), and co-localization with the assembly factor Ktu/Dnaaf2 confirmed that Cfap44 is present in DynAPs (Fig. 4b’). Interestingly, Cfap44 was restricted to only a portion of the DynAPs labelled by Ktu (Fig. 4b’, inset). Because, inner and outer arm dynein subunits are partitioned into sub-regions within DynAPs (Lee et al., 2020), this result is consistent with the role of Cfap44 as a specific tether connecting only a subset of dynein arms (the dimeric, *f*-type inner arms) to axonemal microtubules (Fu et al., 2018; Kubo et al., 2018).

**Figure 4:**
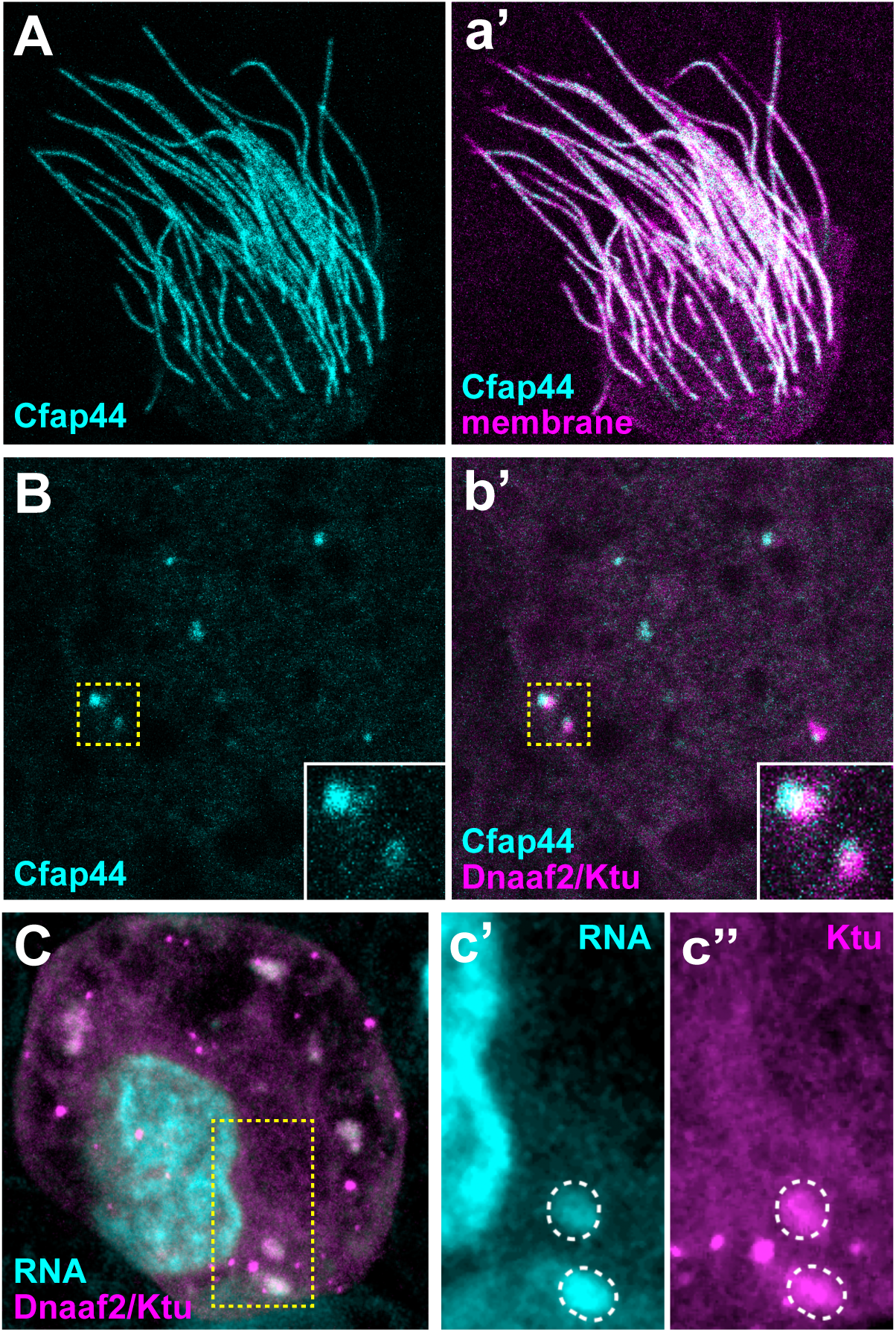
Cfap44 and RNA are present in DynAPs: (A, a’) Cfap44-GFP localizes to axonemes in *Xenopus* motile cilia, as indicated by co-labelling with the membrane-RFP. (B, b’) Cfap44-GFP is also enriched in DynAPs, as indicated by co-labelling with Ktu-GFP. (C, c’, c’’) Staining with CytoRNA Select highlights RNA in the nucleus and also in DynAPs, as indicated by co-labelling with Ktu-RFP.

Finally, RNA-protein interactions are a common mechanism for assembly of liquid-like organelles (Banani et al., 2017; Shin and Brangwynne, 2017), so the localization of Cfap44 prompted us to ask if DynAPs contain RNA. Strikingly, staining of *Xenopus* MCCs with CytoRNA Select revealed both the expected strong signal in nuclei, but also a substantial signal in DynAPs, as indicated by co-localization with Ktu (Fig. 4). Thus, DIF-FRAC identified Cfap44 as an RNA-associated protein, leading to our finding of Cfap44 localization to DynAPs, which in turn led us to discover that RNA is a novel component of DynAPs.

## Discussion

The broad utility of large-scale rosters of RNA-associated proteins is now well-established, yet such resources for vertebrate embryonic tissues remain limited. Thus, both the data and the method reported here will be significant, because they will *a)* provide a valuable resource for understanding mucociliary epithelia and *b)* facilitate future RAP discovery efforts in developing embryos.

The dataset we report provides new hypotheses that should lead to a better understanding of the interaction between microtubule-associated proteins and RNAs generally, as well as specific hypotheses concerning axonemal dynein assembly. Indeed, the only *axonemal* proteins previously found to be present in DynAPs are the dynein subunits themselves (Huizar et al., 2018). Our results therefore suggest the possibility that *f*-type inner arm dyneins may be pre-assembled together with their Cfap43/44 tether in the cytoplasm before deployment to the axoneme. This is in direct contrast to proteins that serve a similar function for the outer dynein arms, such as the “docking complex” proteins Ttc25, Armc4, and Mns1, which are *not* present in DynAPs (Huizar et al., 2018). Moreover, these results may shed light on the etiology of Cfap43/44-related human ciliopathy (e.g. (Coutton et al., 2018; Tang et al., 2017)).

More generally, these data should help us to better understand the connections between ubiquitous liquid-like organelles such as stress granules or P-bodies and cell type-specific organelles such as DynAPs. For example, stress granules and P-bodies share a large number of common components (Aizer et al., 2008; Jain et al., 2016). Likewise, DynAPs are enriched in broadly acting RAPs, such as the heat shock chaperone Hsp90ab1 and the stress granule protein G3Bp1 (Huizar et al., 2018), both of which were identified here as significant DIF-FRAC hits (Supp. Table 1). Likewise, another recent study found that the spliceosome subunit Sf3a3 physically interacts with inner dynein arm subunits and is also enriched in DynAPs (Lee et al., 2020). The data here provide an entry point for a deeper exploration of this interplay between broadly acting RAPs and cell-type specific RAPs such as Cfap44.

Perhaps the largest impact of this work is that it demonstrates the utility of DIF-FRAC for mapping RNA-associated proteomes in embryonic tissues. For example, the method could be readily adopted to explore the mechanisms of polarized mRNA localization in eggs, a ubiquitous developmental mechanism for which *Xenopus* has been a key model (Medioni et al., 2012; Sheets et al., 2017). Our success with animal caps differentiated into mucociliary epithelium also suggests the method will be effective for animal caps experimentally differentiated into any of a wide array of organs and tissues (e.g. (Ariizumi et al., 2009; Okabayashi and Asashima, 2003). Likewise, given the recent interest in studies of explanted tissues from zebrafish (e.g. (Williams and Solnica-Krezel, 2019; Xu et al., 2014), DIF-FRAC could also be readily applied to that powerful animal model.

Ultimately, DIF-FRAC is an entirely label-free method and our use of orthogroups to collapse highly related proteins from multiple sources provides additional power, even when genomes are incompletely annotated or are non-diploid. As such, the findings here open the door to rapid, systematic identification of RNA-associated proteins in *any* model (or non-model) organism for which abundant tissue can be obtained.

## Acknowledgments

This work was supported by grants from the NIH (K99 HD092613 and LRP to K.D), NIH (R01 HL117164 and R01 HD085901) to J.B.W. and/or E.M.M.; R01 DK110520, R35 GM122480 and the Welch Foundation (F-1515) to E.M.M., and a Supplement to Promote Diversity in Health-Related Research from the NICHD (to J.B.W./R.L.H.). Mass spectrometry data collection was supported by CPRIT grant RP110782 to Maria Person and by Army Research Office grant W911NF-12-1-0390.

## Methods

### *Xenopus* animal caps

Xenopus were housed and handled as described (Sive et al., 2000). Animal caps were dissected using forceps and cultured in 1X Steinberg’s solution + gentamicin until sibling embryos reached stage 23 (Nieuwkoop and Faber, 1967). Explants were then lysed for 5 minutes on ice using 500ul PierceIP Lysis Buffer (0.8 mL; 25 mM Tris-HCl pH 7.4, 150 mM NaCl, 1 mM EDTA, 1% NP-40 and 5% glycerol; Thermo Fisher) containing 1x protease inhibitor cocktail III (Calbiochem) and dounced using pestle B. Lysate was then clarified 2x (13,000 g, 10 min, 4 C). Control lysate was left at room temperature for 30 minutes. RNAse lysate was treated with RNAse A (10uL, 100ug, Thermo Fisher, catalog #EN0531) at room temperature for 30 minutes. Lysates were then filtered for multiple rounds (∼5x) (Ultrafree-MC filter unit (Millipore);12,000 g, 2 min, 4 C) to remove insoluble aggregates. The remainder of the DIF-FRAC protocol is as previously described (Mallam et al., 2019) with modifications as discussed in the Results section above.

### Chromatography

Both control and RNAse lysates were subjected to size exclusion chromatography (SEC) independently using a Thermo Scientific UltiMate 3000 HPLC system (Thermo Fisher Scientific, Waltham, MA). Soluble protein (200 μL) was applied to a BioSep-SEC-s4000 600 x 7.8 mm ID, particle diameter 5 μm, pore diameter 500 Å (Phenomenex, Torrance, CA) and equilibrated in PBS, pH 7.2 at a flow rate of 0.5 mL min-1. Fractions were collected every 0.375 mL.

### Mass Spectrometry

Fractions were filter concentrated to 50uL, denatured and reduced in 50% 2,2,2-trifluoroethanol (TFE) and 5 mM tris(2-carboxyethyl)phosphine (TCEP) at 55C for 45 minutes, and alkylated in the dark with iodoacetamide (55 mM, 30 min, RT). Samples were diluted to 5% TFE in 50 mM Tris-HCl, pH 8.0, 2 mM CaCl2, and digested with trypsin (1:50; proteomics grade; 5 h; 37°C). Digestion was quenched (1% formic acid), and the sample volume reduced to 100uL by speed vacuum centrifugation. The sample was desalted on a C18 filter plate (Part No. MFNSC18.10 Glygen Corp, Columbia, MD), eluted, reduced to near dryness by speed vacuum centrifugation, and resuspended in 5% acetonitrile/ 0.1% formic acid for analysis by LC-MS/MS. Peptides were separated on a 75uM × 25 cm Acclaim PepMap100 C-18 column (Thermo) using a 3%–45% acetonitrile gradient over 60 min and analyzed online by nanoelectrospray-ionization tandem mass spectrometry on an Orbitrap Fusion or Orbitrap FusionLumos Tribrid (Thermo Scientific). Data-dependent acquisition was activated, with parent ion (MS1) scans collected at high resolution (120,000). Ions with +2 or higher charge were selected for HCD fragmentation spectrum acquisition (MS2) in the ion trap, using a Top Speed acquisition time of 3 s. Dynamic exclusion was activated, with a 60 sec exclusion time for ions selected more than once. MS was acquired in the UT Austin Proteomics Facility.

### Reference Proteome Construction

Two *Xenopus laevis* protein database were downloaded from Xenbase (http://www.xenbase.org/) (Karpinka JB et al., 2015; Nenni et al., 2019): the *X. laevis* JGI gene model (v9.1) peptide FASTA (http://ftp.xenbase.org/pub/Genomics/JGI/Xenla9.1/1.8.3.2/XL_9.1_v1.8.3.2.primaryTranscripts.pep.fa.gz) and an *X. laevis* protein sequence file containing GenBank sequences (http://ftp.xenbase.org/pub/Genomics/Sequences/xlaevisProtein.fasta). The files were filtered to remove sequences shorter than 20 amino acids or containing greater than 30% X’s (*i.e.*, nonstandard amino acid symbol). A combined proteome was made by concatenating the JGI gene model FASTA (XL_9.1_v1.8.3.2.primaryTranscripts.pep.fa) and the GenBank sequence FASTA (xlaevisProtein.fasta). Each sequence in the combined file was mapped to EggNOG orthogroups using emapper (v 0.12.7) and vertebrate-level Hidden Markov models downloaded from the eggNOG database in August 2017. An orthology-collapsed proteome was then constructed by grouping protein sequences based on shared EggNOG orthology groups. Sequences were then concatenated separating each protein’s sequence by triple lysines to ensure theoretical tryptic cleavage sites even when considering up to two missed tryptic cleavages. Proteins that did not map to any vertebrate-level eggNOG groups were retained in their original FASTA entry format.

Since *X. laevis* homeologs and near-duplicate Xenbase/Genbank protein entries were combined into a single protein entries *via* the triple lysine concatenation, the orthology collapsed proteome is usable by any MS/MS search engine, which then processes the concatenated proteins properly into peptides (but for a single tailing lysine on the C-terminal-most peptide of internal proteins in the concatenation). Most significantly, by combining high sequence similarity proteins, the collapsed proteome substantially increases the rate of unique peptide assignment and overall protein identification (Supplemental Table 1). After database spectral matching, entries described by eggNOG groups are related back to individual proteins *via* an annotation table mapping eggNOG IDs to eggNOG annotations, eggNOG IDs to Xenbase/Genbank IDs and XenBase/GenBank annotations, and eggNOG IDs to human UniProt IDs and human UniProt annotations.

The orthology collapsed proteome (XL9.1_XLgb_concat_orthocollapse_01102020.fasta) as well as accompanied scripts can be downloaded here: https://github.com/marcottelab/pivo.

### Protein identification

Raw formatted mass spectrometry files were first converted to mzXML file format using MSConvert (http://proteowizard.sourceforge.net/tools.shtml) and then processed using the MSBlender protein identification pipeline (Kwon et al., 2011), combining peptide-spectral matching scores from MSGF+ (Kim and Pevzner, 2014), X! TANDEM (Craig and Beavis, 2004) and Comet (Lingner et al., 2011) as peptide search engines with default settings. A false discovery rate of 1% was used for peptide identification. Elution profiles were assembled using unique peptide spectral matches for each EggNOG orthogroup across all fractions collected.

### DIF-FRAC Analysis

To identify proteins that are sensitive to RNAse treatment, we used our previously described statistical framework (Mallam et al., 2019). Briefly, each protein’s control and RNAse elution profiles (Supp. Tables 4-7) are compared using a normalized Manhattan distance called the DIF-FRAC score to measure the overall change between profiles. Next, for each protein, we calculate a background distribution of DIF-FRAC scores. The background distribution is created from proteins with similar overall mass spec observed abundances to the target protein. More specifically, the distribution is made of a window of proteins with abundance rank +100 and −100 surrounding the target protein. Additionally, proteins with known RNA binding annotations are removed from the background distribution. The distribution is then modeled using a two-component Guassian mixture model where the component with the highest-mean represents RNA associated proteins, and the component with the lowest-mean represents non-RNA associated proteins (*i.e.*, background distribution). We then calculate a Z-score by comparing the DIF-FRAC score of the target protein to the background distribution. A *p*-value is calculated from the Z-score and finally, the *p*-value is false discovery corrected using Benjamini/Hochberg correction (Benjamini and Hochberg, 1995).

### Systems-level analyses

For system-wide analyses in Figure 1, eggNOG entries with significant DIF-FRAC scores (DIF-FRAC *p*-value < 0.05) were mapped to human Uniprot accessions. The gProfiler website (Raudvere et al., 2019) was used for Gene Ontology molecular function enrichment. Background proteins were defined to be the set of all identified proteins in the DIF-FRAC experiment. Electronic annotations were not considered in the analysis. Default values were used for all other parameters. For the Interpro superfamily enrichment analysis, Uniprot accessions were mapped to Interpro superfamily ids (Hunter et al., 2009) via Uniprot’s ID mapping web service. Background distributions were calculated by taking random samples from all identified eggNOG entries. The occurrence of each Interpro superfamily was tabulated for each random sample and the mean and standard deviation were calculated across 1000 total random samplings.

### Protein Structure Modelling

To model the structure of Cfap44, we used a template-based homology modeling approach. We first identified suitable templates for Cfap44 using default parameters from pGenTHREADER web server tool (Lobley et al., 2009) targeting amino acid sequence from position 400 to 1600 in Xenopus Cfap44 protein. We then used MODELLER (Sali and Blundell, 1993) to build the structure of Cfap44 using PDB ID: 2YMU as a template. The alignment of the Cfap44 structure model with Gemin5 (PDB ID: 5GXH) was done using Chimera (Pettersen et al., 2004).

### Imaging

Full length of *Xenopus cfap44* was identified from Xenbase (Nenni et al., 2019), was amplified from *Xenopus*cDNA library and was cloned into pCS10R vector fused with N-terminal GFP. Capped cfap44 mRNA was synthesized using mMESSAGE mMACHINE SP6 transcription kit (ThermoFisher Scientific). Each 90 pg of GFP-cfap44 and mCherry-Dnaaf2/Ktu mRNAs were injected into two ventral blastomeres and live-imaging was performed as previously described (in Huizar et al., 2018). For RNA staining, embryos were fixed with Dent’s fixative (20% DMSO and 80% methanol) at stage 23 and were stained by 500 nM of SYTO RNAselect Green Fluorescent Cell Stain (Invitrogen) for 20min and were imaged after washing.

### Data Deposition

Proteomics data has been deposited in the PRIDE repository with accessions PXD017659 and PXD017650.

### Code Repository

Source code is freely available on GitHub: https://github.com/marcottelab/diffrac

---

**Supplemental Figure 1:**
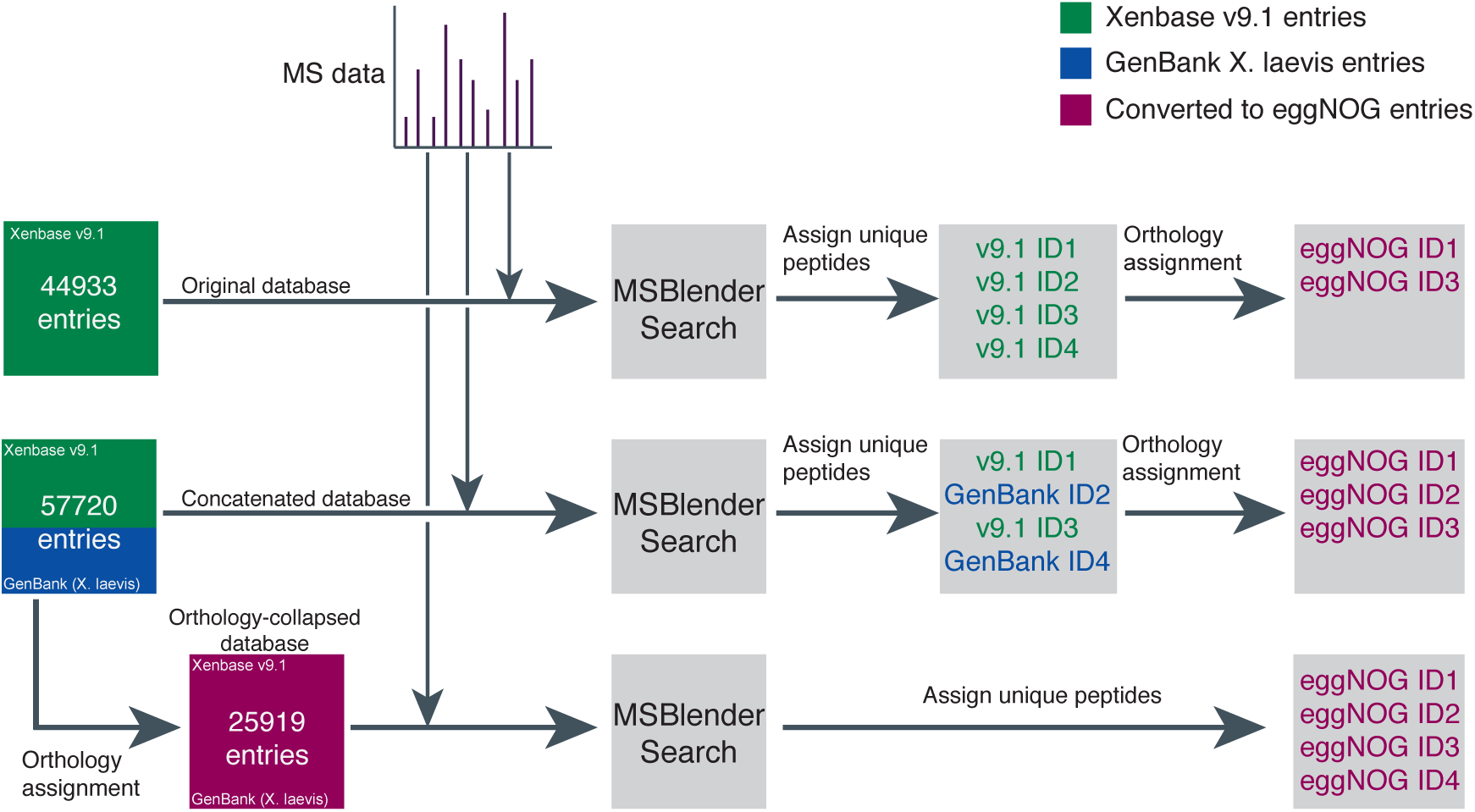
Flowchart indicating steps taken to directly compare different versions of the *X. laevis* proteome. Three proteomes were compared 1) an unaltered database obtained from the v9.1 genome assembly (green) 2) a combination of v9.1 proteins with GenBank proteins (green and blue) and 3) an EggNOG ortho-collapsed proteome derived from the combination of v9.1 and GenBank. Mass spectra from the control size exclusion experiment was analyzed using MSBlender and the three proteomes as reference proteomes. To compare the orthology-collapsed proteome to the un-collapsed proteomes identified proteins from the uncollapsed MSBlender searches were mapped *post hoc* to EggNOG orthology groups. The final performance evaluation of each was evaluated based on the number of orthology groups identified (Supplementary Table 1).

**Supplemental Figure 2:**
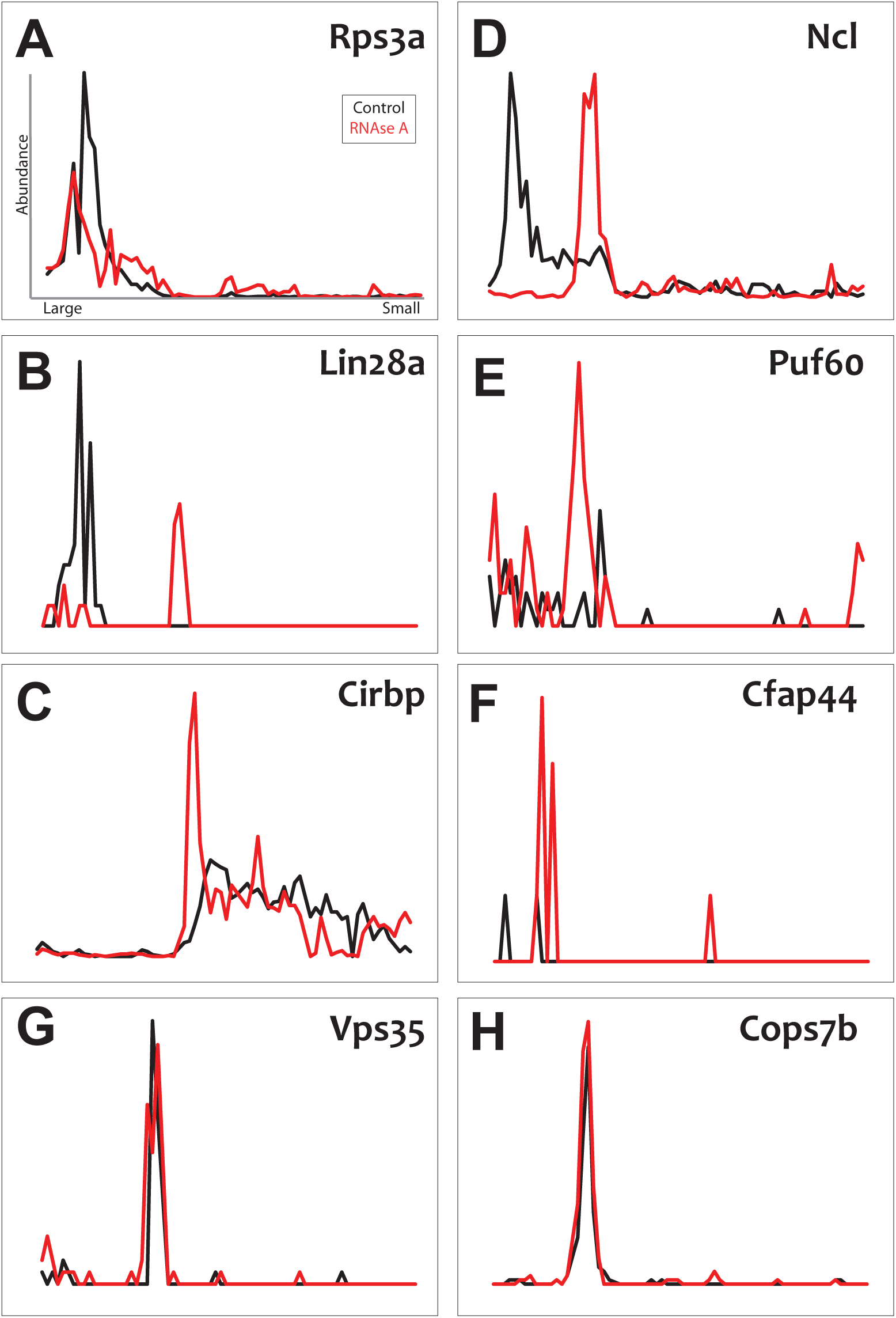
Elution profiles from a biological replicate RNAse DIF-FRAC experiment shows consistency with replicate 1. (A-E) Replicate elution profiles of known RNA-binding proteins show consistent shift upon RNAse treatment. (F) Replicate elution profile of Cfap44, a ciliopathy gene identified as RNA associated. (G-H) Replicate elution profiles of negative controls which do not show shift upon RNAse treatment.

**Supplemental Figure 3:**
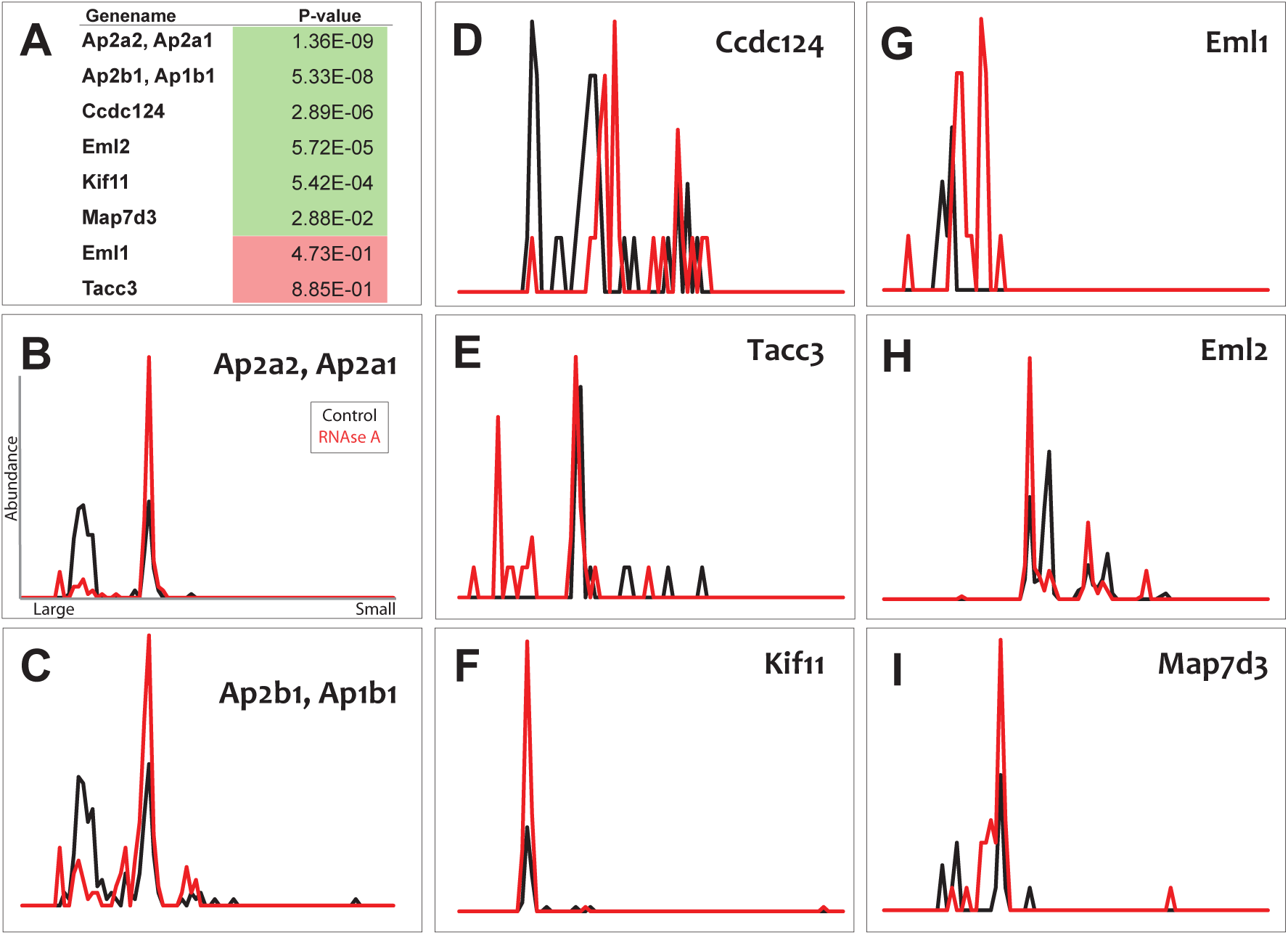
Elution profiles of AP-2 complex subunits and microtubule-associated proteins. (A) Table of DIF-FRAC calculated *p*-values for AP-2 complex subunits and microtubule-associated proteins. (B-C) AP-2 complex subunits elution profiles show a shift upon RNAse treatment. (D-I) Elution profiles of microtubule-associated proteins show shifts upon RNAse treatment.

**Supplemental Table 1:** Comparison of proteomes used in the mass spectrometry pipeline. Protein and orthogroup identifications are made based on unique peptide matching to ungrouped entries. To compute the number of identifications in a way directly comparable to the orthology-collapsed proteome, the protein identifications made by the first two proteomes were *post hoc* assigned to eggNOG groups (Supplementary Figure 1). Thus, the total number of IDs in this table are sourced from the same pool of eggNOG-assigned groups of proteins (in addition to individual protein entries that do not map to any vertebrate-level orthology group) and can be directly compared to one another. Unique IDs represent proteins found only when that particular proteome is searched against, i.e., the orthology-collapsed proteome nets a significant amount of information that is inaccessible when using the other un-collapsed proteomes.

| Proteome | PSMs > 1 |  | PSMs > 10 |  |
| --- | --- | --- | --- | --- |
|  | Total #<br>OrthoIDs* | Unique<br>OrthoIDs* | Total #<br>OrthoIDs* | Unique<br>OrthoIDs* |
| v9.1 sequences | 5244 | 221 | 1770 | 40 |
| v9.1 sequences + GenBank<br>sequences | 3127 | 49 | 710 | 1 |
| v9.1 sequences + GenBank<br>sequences (orthology collapsed) | 6391 | 1131 | 2371 | 563 |
| Wuhr transcriptome | 3630 | 412 | 1163 | 37 |

**Supplemental Table 2: DIF-FRAC result table with FDR adjusted *p*-values.**

**Supplemental Table 3: Biological replicate DIF-FRAC result table with FDR adjusted *p*-values.**

**Supplemental Table 4: Elution matrix of control experiment (replicate 1).** Tables 4-7 are text files containing elution data, which we designate as “.elut” files. Values in entries are peptide spectral matches. Columns designate fractions; rows designate proteins/orthogroups. These files can be used with our DIF-FRAC software (deposited in GitHub, see Methods, below) to generate graphical elution profiles.

**Supplemental Table 5: Elution matrix of rnaseA experiment (replicate 1).**

**Supplemental Table 6: Elution matrix of control experiment (replicate 2).**

**Supplemental Table 7: Elution matrix of rnaseA experiment (replicate 2).**

